# RSGSA: a Robust and Stable Gene Selection Algorithm

**DOI:** 10.1101/2020.07.27.216879

**Authors:** Subrata Saha, Ahmed Soliman, Sanguthevar Rajasekaran

## Abstract

Nowadays we are observing an explosion of gene expression data with phenotypes. It enables researchers to efficiently identify genes responsible for certain medical condition as well as classify them for drug target. Like any other phenotype data in medical domain, gene expression data with phenotypes also suffers from being very underdetermined system. In a very large set of features but a very small sample size domains (e.g., DNA microarray, RNA-seq data, GWAS data, etc.), it is often reported that several different spurious feature subsets may yield equally optimal results. This phenomenon is known as *instability*. Considering these facts, we have developed a very robust and stable supervised gene selection algorithm to select the most discriminating non-spurious set of genes from the gene expression datasets with phenotypes. *Stability* and *robustness* is ensured by class and instance levels perturbations, respectively.

We have performed rigorous experimental evaluations using 10 real gene expression microarray datasets with phenotypes. It revealed that our algorithm outperforms the state-of-the-art algorithms with respect to stability and classification accuracy. We have also done biological enrichment analysis based on gene ontology-biological processes (GO-BP) terms, disease ontology (DO) terms, and biological pathways.

## 1 Introduction

Gene expression is the process by which information from a gene is used in the synthesis of a functional gene product such as proteins. The genetic code preserved in a DNA sequence is “interpreted” by gene expression, and the properties of the expression give rise to the organism’s phenotypes (i.e., observable traits such as presence of a disease, height, etc.). Therefore, regulation of gene expression is critical to an organism’s development. In the field of molecular biology gene expression profiling is the measurement of the activity of thousands of genes simultaneously to create a global picture of cellular function. Several transcriptomics technologies have been developed to produce necessary data to analyze. DNA microarrays measure the relative activity of previously identified target genes. Sequence based techniques, like RNA Sequencing (RNA-seq), provide information on the sequences of genes in addition to their expression level. RNA-seq based on next-generation sequencing (NGS) technology enables transcriptome analyses of entire genomes at a very high level of resolution. These procedures are not only very cost effective but also can be done in laboratory environment.

Like any other phenotype data in medical domain, gene expression data consisting of phenotypes also suffer from being very underdetermined system. In other words, for most of the gene expression data the number of samples are very small compared to the large number of genes or transcripts. As the quality of genome sequences and the methods for identifying protein-coding genes improved [16], the count of recognized protein-coding genes dropped to 19, 000 – 20, 000 [11]. However, a complete understanding of the role played by genes expressing regulatory RNAs that do not encode proteins has raised the total number of genes to at least 46, 831 [30] plus another set of 2, 300 micro-RNA genes [1]. Since the number of samples varies from few hundreds to few thousands, gene selection is particularly challenging in a very underdetermined system as the system can have many spurious solutions. The algorithm of choice may select the random genes instead of the real set of discriminating genes.

Traditional statistical methods [3] are designed to analyze susceptibility of genes from gene expression data with phenotype by considering only a single gene at a time. On the contrary, it is proven that multiple genes act together to cause many common diseases. There are numerous challenges in designing and analyzing joint effects of multiple differentially expressed genes. Nowadays with next generation sequencing methods (e.g., RNA-seq, CAGE, etc.) specific transcript expression can be identified. The total number of human transcripts with protein-coding potential is estimated to be at least 204,950 [15]. As stated earlier the expressions of those transcripts are measured from not more than several thousands individuals. Here we have many more variables *p* than samples *n*. Classical model requires *p* < *n* where we have *p* ≫ *n*. As a consequence, standard multi-variable statistical approaches are not very promising tools to detect complex gene-gene interactions from genome-wide data. On the contrary machine learning algorithms provide several intuitive alternatives to perform multi-gene analyses within acceptable time, accuracy, memory, and money.

However, machine learning tools also suffer from very underdetermined systems where we have many more variables compared to the number of samples. Fortunately, ensemble technique can be employed in the context of supervised gene selection domain. In statistics and machine learning, ensemble learning is a machine learning technique where multiple learners are trained to solve the same problem. In contrast to ordinary machine learning approaches which try to learn one hypothesis from training data, ensemble methods try to construct a set of hypotheses and combine them to use. We could obtain better predictive performance by combining multiple learning algorithms instead of employing only one learner [14]. Specifically, generalization ability of an ensemble is usually much stronger than that of a single learner [35]. In this article we have proposed an ensemble gene selection algorithm dubbed as “Robust and Stable Gene Selection Algorithm” (RSGSA, in short) that is stable and robust compared to other state-of-the-art algorithms.

## 2 Methods

### 2.1 Approach

Feature selection is the problem of identifying a subset of the most relevant features in the context of model construction. If the total number of features is *n*, the total number of candidate subsets will be 2^*n*^. An exhaustive search strategy searches through all the 2^*n*^ feature subsets to find an optimal one. Clearly, this may not be feasible in practice [20]. A number of heuristic search strategies have been introduced to overcome this problem. Subset selection begins with an initial subset that could be empty, the entire set of features, or some randomly chosen features. This initial subset can be changed in a number of ways. In forward selection strategy, features are added one at a time. In backward selection the least important feature is removed based on some evaluation criterion. Random search strategy randomly adds or removes features to avoid being trapped in a local maximum.

Depending on the type of data, feature selection can be classified as supervised, semi-supervised, and unsupervised. A data instance (e.g., a patient potentially having cancer) is characterized by a number of independent variables (features), e.g., tumor markers (substances found in the blood, urine, stool, other bodily fluids, or tissues of the patient). It may also have a response variable (often called a label), e.g., whether the patient has a benign or a malignant tumor. If all the data instances in the data set have known response values, the process of feature selection is called “supervised”. Supervised feature selection techniques can be broadly classified into 3 categories: (1) wrapper, (2) filter, and (3) embedded.

#### Wrapper

In a wrapper method the classification or prediction accuracy of an inductive learning algorithm of interest is used for evaluation of the generated subset. For each generated feature subset, wrappers evaluate its accuracy by applying the learning algorithm using the features residing in the subset. Although it is a computationally expensive procedure, wrappers can find the subsets from the feature space with a high accuracy because the features match well with the learning algorithm [21]. Examples include evolutionary algorithms [39], simulated annealing [23], and randomized search [32].

#### Filter

Filter methods are computationally more efficient than wrapper methods since they evaluate the accuracy of a subset of features using objective criteria that can be tested quickly. Common objective criteria include the mutual information, Pearson product-moment correlation coefficient, and the inter/intra class distance. Though filters are computationally more efficient than wrappers, often they produce a feature subset which is not matched with a specific type of predictive model and thus can yield worse prediction accuracies. Some notable methods include symmetric uncertainty (SU) [29], gain ratio (GR) [17], Kullback-Leibler divergence measure (KLD) [36], and RELIEF [22].

#### Embedded

Embedded methods combine the qualities of filter and wrapper methods. It’s implemented by algorithms that have their own built-in feature selection methods. Some of the most popular examples of these methods are linear support vector machine (LSVM) [13], random forest (RF) [7], least absolute shrinkage and selection operator (LASSO) [34], and ridge regression [38] which have inbuilt penalization functions to reduce overfitting.

If only some data instances have known response values, we are facing a semi-supervised feature selection problem. If none of the data instances have response values, it is “unsupervised” feature selection. The majority of research efforts is in the area of supervised feature selection.

As stated earlier, ensemble gene selection techniques might be employed in the domain of supervised gene selection algorithms to ensure *stability* and *robustness*. In this context we define the *stability* and *robustness* as they have some clear distinction between them. In the context of gene selection algorithm we can define *robustness* as the variation of selected genes resulting from small changes in the dataset such as adding or removing samples from the dataset. Like-wise, We can define the stability of gene selection algorithms as the variation in gene selection results due to adding small noise in the dataset. The less the variation the more the algorithm will be robust and stable. Both of robustness and stability are desirable characteristics of a gene selection algorithm, especially where the number of biomarkers are ways more than the number of samples. To ensure *robustness* we bootstrapped the samples multiple times to create slightly smaller datasets by randomly taking samples with replacement. For each dataset we run our gene ranking algorithm. To ensure stable ranking a small amount of noise is introduced in the ranking algorithm by randomly flipping class labels of a few samples. To get in-depth knowledge about ensemble techniques for feature selection readers are referred to [5]. To the best of our knowledge, the algorithmic steps we follow such as, removing correlated features and ensuring stability in SVM-RFE’s recursive stage are unique in this domain. Next we describe our algorithmic framework in detail.

### 2.2 Removing correlated genes

In general, correlated genes do not improve the learning model of interest. There are basically 3 main reasons to remove correlated genes from the set of given genes: (1) making the gene selection algorithm faster; (2) decreasing harmful bias; and (3) making the model simpler and interpretable.

We have designed and developed a novel graph theoretic algorithm to effectively remove the correlated genes from consideration. It works as follows: Suppose we have a set of genes *S*. A graph *G* is constructed in which there is a node for each gene *s* ∈ *S*. Two nodes *n*_1_ and *n*_2_ in *G* will be connected by an edge *e* if *r*(*n*_1_, *n*_2_) ≥ *λ* where *r* is the Pearson’s correlation coefficient and *λ* is a user defined threshold. In our experiments we set *λ* = 0.9. After constructing such a graph *G*, we extract all the connected components of *G*. For each connected component we measure *eigenvector centrality* [25] of each node residing in that component. In graph theory, *eigenvector centrality* (also called *eigencentrality*) is a measure of the influence of a node in a network. Relative scores are assigned to all nodes in the network based on the concept that connections to high-scoring nodes contribute more to the score of the node in question than equal connections to low-scoring nodes. A high eigenvector score means that a node is connected to many nodes who themselves have high scores. Let a node *n* have the highest eigenvector score across all the connected components. We delete all the neighboring nodes of *n* and *n* from *G*. We record *n* as a *leader* of its neighbors. Since all the neighbors of *n* are connected with *n*, they are highly correlated with *n*. So, deleting the neighbors of *n* will not only cost minimal information loss but also reduce the dimension of the gene space. The same procedure is repeated until all the nodes *n* ∈ *G* are isolated i.e., there is no edge *e* in *G*. We record all such nodes *n* as *leaders*. The details of the algorithm can be found in Algorithm 1.

#### Algorithm 1

**R**emove **C**orrelated **G**enes (RCG)

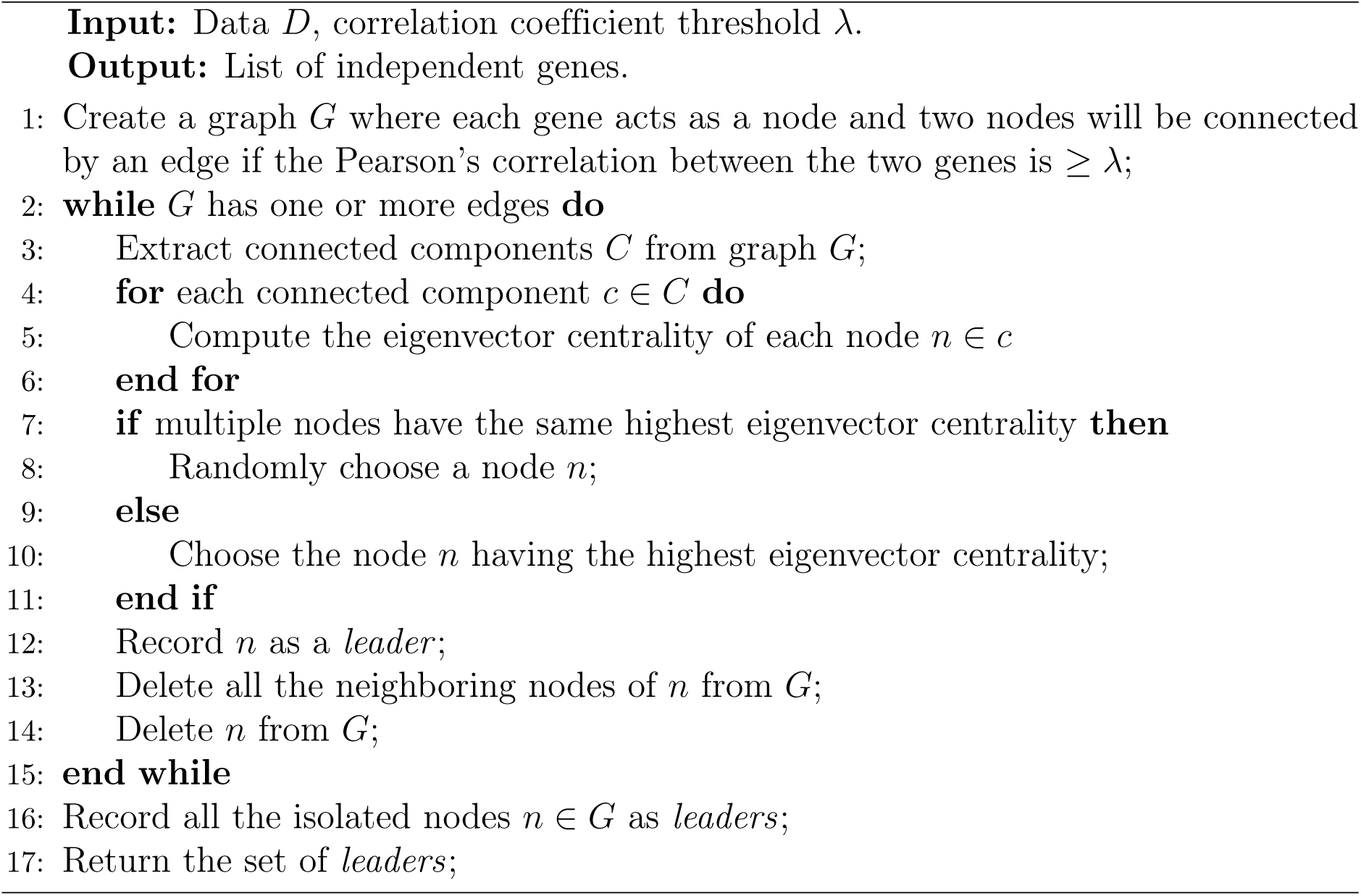

#### Algorithm 2

**R**obust and **S**table **G**ene **S**election **A**lgorithm (RSGSA)

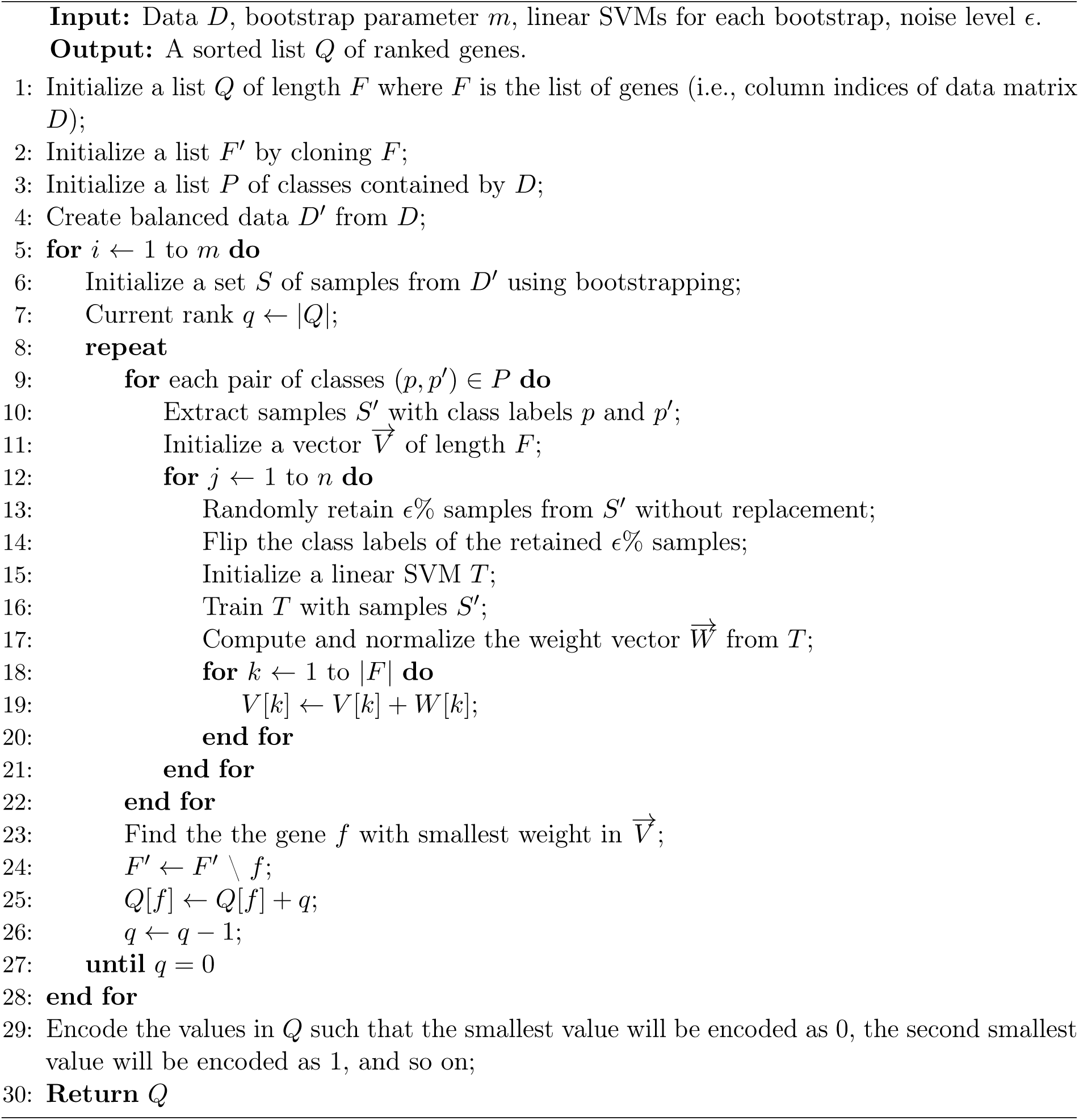

### 2.3 Ranking genes

Recursive feature elimination method based on support vector machines (SVM-RFE, in short) was first proposed in [13] to rank genes based on their importance for cancer classification. It works by recursively removing one or more weak features (e.g., genes) until the specified number of features is reached. At the beginning, a linear SVM is trained on the initial set of features to build an inductive model. In binary classification problem this model is nothing but a vector of weights (also known as coefficient vector) describing a separating hyperplane between two classes. Each entry of the vector corresponds to the coefficient of a particular feature. The importance of each feature is directly proportionate with its coefficient in the weight vector [13]. The more the weight of a feature the more will it be important. The least important features are pruned from the current set of features using the weight vector. This procedure is recursively repeated on the current set of features until the desired number of most important features is eventually reached. We attempt to make SVM-RFE stable by introducing small changes in class labels and employing ensembles of SVMs in each recursive step to eliminate a set of least important genes. Due to this small change, the weight associated with each gene will not be identical for the identical dataset in different runs. By averaging the weights across a set of weight vectors from linear SVMs of the same configuration, we can make the weight vector stable. Since weights are directly associated with the importance of the genes, ranking should also be stable. We call this variation of SVM-RFE as Stable SVM-RFE (SSVM-RFE, in short). Let the number of recursive steps taken by SSVM-RFE to rank a set *S* of genes of interest be *R*. Suppose in each of the *R* recursive steps we employ a set *L* of linear SVMs. Each SVM *l* ∈ *L* is trained to build an inductive model by introducing small changes in class labels and we get a corresponding weight vector *w*_*l*_ (1 ≤*l* ≤ |*L*|). According to [13] the importance of the *i*^*th*^ gene 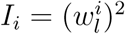 where 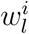 represents the *i*^*th*^ weight component of 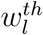 weight vector. At the end of each recursive step we get a set of |*L*| weight vectors. Let the set of weight vectors obtained in the *r*th recursive step be denoted as *W*_*r*_, 1 ≤ *r* ≤ *R*. Since in each run and model we introduce some small noise in class labels, the weight vectors will be different from each other. To make it stable we normalize each weight vector and average each component. Let the number of genes remaining in a particular run be *n*. Each weight vector *w*_*l*_ will have *n* components and is normalized as follows:

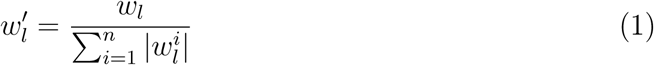

The *i*^*th*^ component of the final weight vector *W*_*r*_ for the *r*th recursive step (1≤ *r* ≤ *R*) is formed as follows:

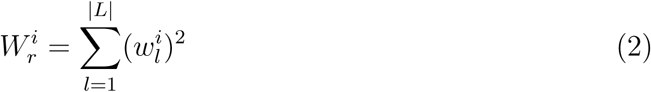

At the end of the *r*th recursive step, the importance of the *i*^*th*^ gene is defined as 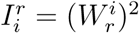 and we discard one or more genes having the least scores. The above procedure is done recursively until we have the desired number of genes left. Consequently, SSVM-RFE outputs the most important genes. We claim that these ranks are stable.

To make our algorithm robust we repeatedly run the above procedure on different bootstrapped samples. If the dataset is imbalanced, our algorithm automatically makes it balanced before bootstrapped sampling by creating synthetic samples from the minority class instead of creating copies [10]. The final ranking is done by aggregating all the rankings produced from different samples. Let the number of SSVM-RFE runs employed be *m*. Each SSVM-RFE run produces a list of stable ranks for the given set of genes *S*. Let the *i*^*th*^ SSVM-RFE run provide a gene ranking 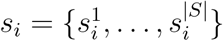 where 1 ≤ *i* ≤ *m*. We can aggregate the gene rankings by taking the sums as follows:

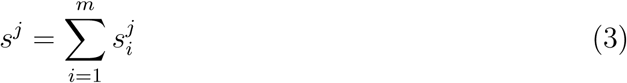

where *j* represents the *j*^*th*^ gene. The pseudo code of our algorithm can be found in Algorithm 2. Please, note that the input data matrix *D* is formed using Algorithm 1.

### 2.4 Analysis

In this section we analyze the time complexity of the algorithm. Let *n* be the number of genes in the input. Let *e* be the number of training examples. In Algorithm 1, we can construct the graph *G*(*V, E*) in time *O*(*n*^2^*e*), since the Pearson’s correlation coefficient between any two genes can be computed in *O*(*e*) time. Given a graph *G*(*V, E*), the problem of computing the eigencentrality for each node can be reduced to the problem of computing the eigen vectors of a |*V*| × |*V*| matrix [9]. As a result, if there are *n* genes, then their eigencentrality values can be computed in *O*(*n*^3^) time (since eigen vectors for a *n* ×*n* matrix can be computed in *O*(*n*^3^) time [27, 28]).

In Algorithm 1, we start with *n* genes, find a node with the highest eigencentrality, and remove all of its neighbors and itself. We repeat the process of isolating a node and its neighbors until there are no more edges in the graph. This process is described in lines 2 to 14. In each iteration of this while loop the number of remaining genes decreases. Note that in the worst case only one edge (and two nodes) from the graph might get deleted. This means that the total time for computing the eigencentrality for the nodes across all the iterations of the while loop is *O*(*n*^4^). Please note that this is the worst case which does not arise in practice. In practice, our algorithm is very fast.

Thus the run time of Algorithm 1 is *O*(*n*^2^*e* + *n*^4^).

In Algorithm 2, let *N* be the starting number of genes. For example, let *N/*2 be the target number of genes to be output. In each iteration of the repeat loop of line 8, a constant fraction of the genes with the lowest weights are eliminated. In other words, the repeat loop is executed only *R* = *O*(1) times. For instance, in the first iteration we eliminate ^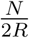^ genes; In the second iteration we eliminate another set of ^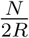^ genes, and so on.

In every iteration of the repeat loop we have to train a linear SVM |*L*| times. The run time of linear SVM is *O*(*ab* min{*a, b*}) where *a* is the number of attributes and *b* is the number of training examples (see e.g., [18]). As a result, the time spent in one execution of the repeat loop is *O*(*L RNe* min *N, e*). The for loop of line 5 is executed *m* times. As a result, the total run time of Algorithm 2 is *O*(*m*|*L*|*RNe* min{*N, e*}).

Put together, the run time of our algorithm is *O*(*n*^2^*e* + *n*^4^ + *m*|*L*|*RNe* min{*N, e*}). If *n > e*, this run time will be *O*(*n*^4^ + *m*|*L*|*Rne*^2^). On the other hand, if *e* ≥ *n*, then the run time will be *O*(*n*^2^*e* + *n*^4^ + *m*|*L*|*Rn*^2^*e*). We arrive at the following Theorem:

**Theorem 2.1** *The run time of RSGSA is O*(*n*^2^*e* + *n*^4^ + *m*|*L*|*RNe* min{*N, e*}). *If n > e, this run time will be O*(*n*^4^ + *m*|*L*|*Rne*^2^). *On the other hand, if e* ≥ *n, then the run time will be O*(*n*^2^*e* + *n*^4^ + *m*|*L*|*Rn*^2^*e*).

## 3 Results

### 3.1 Datasets

To demonstrate the effectiveness of our algorithm RSGSA, we have used 10 real microarray gene expression datasets. A microarray is a laboratory tool used to detect the expression of thousands of genes simultaneously where each expression of a gene is a real valued number. Each row of a gene expression dataset corresponds to an individual where each column represents the expression of a particular gene. Details of the datasets used can be found in Table 1. More details about the datasets can be found in [43]. It is to be noted that our algorithm works on any kind of gene expression datasets with phenotypes such as, RNA-Seq data.

**Table 1.**
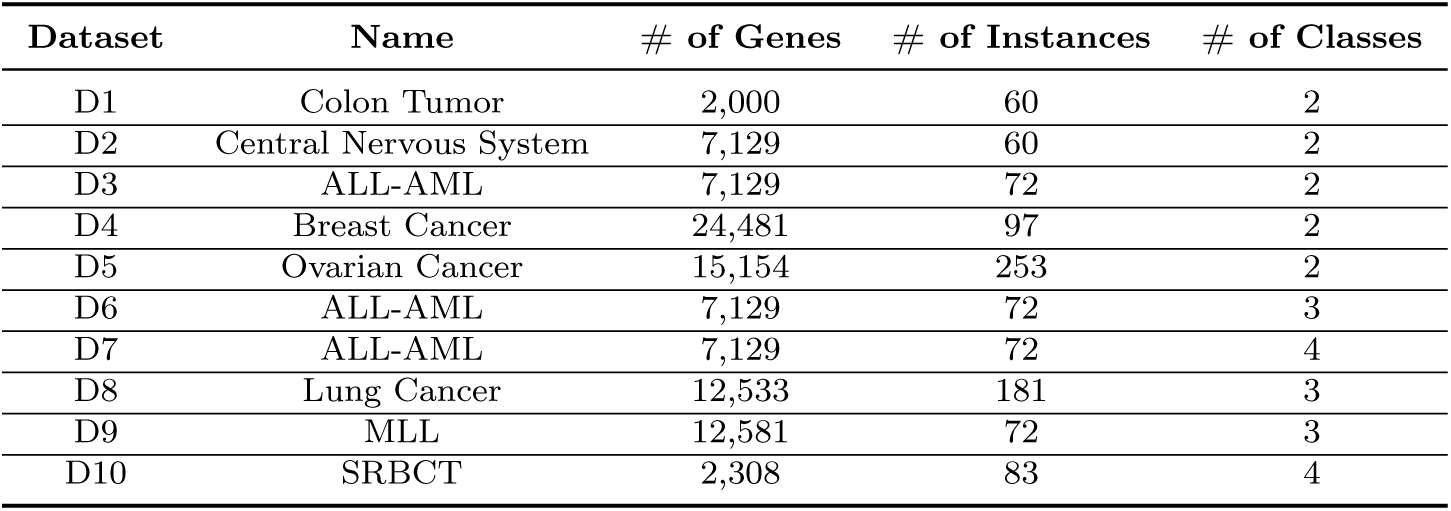
Datasets used for gene selection

### 3.2 Experimental setup

All the experiments were done on an Intel Westmere compute node with 12 Intel Xeon X5650 Westmere cores and 48 GB of RAM. The operating system running was Red Hat Enterprise Linux Server release 5.7 (Tikanga). The algorithm is written in standard Java programming language. Java source code is compiled and run by Java Virtual Machine (JVM) 1.8.0.

### 3.3 Evaluation metrics

We measure the effectiveness of our proposed algorithm RSGSA using 3 different performance metrics. These metrics are defined below.

#### 3.3.1 Jaccard index

*Stability* is measured by employing Jaccard index. The Jaccard index, also known as Intersection over Union and the Jaccard similarity coefficient, is a statistic used for gauging the similarity and diversity of sample sets. The Jaccard coefficient measures similarity between finite sample sets, and is defined as the size of the intersection divided by the size of the union of the sample sets: 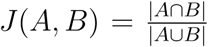 where *A* and *B* are two sample sets.

The Jaccard index varies from 0.0 to 1.0. The higher the value, the more similar will be the two sets. Although it’s easy to interpret, it is extremely sensitive to small samples sizes and may give erroneous results, especially with very small samples or data sets with missing observations.

#### 3.3.2 Informedness

We are interested in evaluating the goodness of a classifier *C* in correctly identifying positive and negative examples from a set of examples. For instance, positive and negative examples in binary classification problem can be referred to as persons having a specific disease and healthy individuals, respectively. We refer to positive examples that are identified as positive as true positives (TP), positive examples that are identified as negative as false negatives (FN), negative examples that are identified as positive as false positives (FP), and negative examples that are identified as negative as true negatives (TN) by a classification algorithm *C*.

*Informedness* is a measure of how informed system *C* is about positives and negatives, i.e., 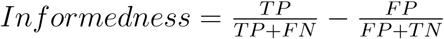. More higher the value, more higher will be the *informedness*. It varies from −1.0 to +1.0. A value of +1 implies *C* is fully informed about positives and negatives. Since *informedness* is dependent on all of these numbers (e.g., TP, FP, TN, and FN), they are sensitive to any change to (the proportions of) these numbers. Therefore, when both positives and negatives are of interest, *informedness* is more informative evaluation measures for a classification algorithm *C*. In this article *informedness* is also termed as *accuracy* i.e., these two terms are interchangeable throughout this article. Note that *informedness* is measured using 10-*fold* cross validation.

#### 3.3.3 Gain

Gain measures the percentage improvement over any performance metric (such as Jaccard index, informedness, etc.) achieved by RSGSA when compared to other algorithms. Let the performance metric of interest of RSGSA and other algorithm of interest be *p*′ and *p*′′, respectively. The gain is measured using this formulation: 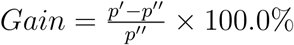.

### 3.4 Outcome

As noted above we have done performance evaluation using 10 real gene expression microarray datasets. 5 feature selection algorithms (namely, symmetric uncertainty (SU), gain ratio (GR), Kullback-Leibler divergence (KLD), RELIEF, and SVM-RFE) were used to evaluate the performance of our algorithm RSGSA. We have also employed random forest (RF) based attribute evaluation method but we are not reporting its consistent poor performance over all the datasets. Suppose a particular algorithm and dataset be *A* and *D*, respectively. For each dataset *D* we execute each algorithm *A* 10× ; each time we randomly chose 80% samples from *D* with replacement. After each execution of *A* we record top *X* genes where *X* = {50, 100, 150, 200}. We then compute pair-wise Jaccard indices for each of *X* genes and take the average over all the pair-wise indices. The procedure was done for all the dataset *D* by running each method *A*.

In a similar fashion we compute the classification accuracy (i.e., *informedness*) by taking top *X* genes selected by the algorithms. It woks as follows: We extract dataset *D*′ containing only top *X* genes and their corresponding expression values across the samples. We execute classifier LSVM 10 × ; each time we randomly chose 80% samples from *D*′ with replacement. After each execution of LSVM we record classification accuracy (i.e., *informedness*) for top *X* genes where *X* = {50, 100, 150, 200}. We then compute pair-wise classification accuracy for each of *X* genes and take the average over all the pair-wise accuracy. The classification accuracy is measured using 10-*fold* cross validation. Let us assume *D*′ is matrix were each row refers to a sample (i.e., individuals) and each column contains the gene expression value of a particular gene from top *X* genes. In 10-*fold* cross-validation, the original sample (*D*′) is randomly partitioned into 10 equal sized subsamples. Of the 10 subsamples, a single subsample is retained as the validation data for testing the model, and the remaining 9 subsamples are used as training data. The cross-validation process is then repeated 10 times, with each of the 10 subsamples used exactly once as the validation data. The 10 results (i.e, *informedness*) can then be averaged to produce a single estimation. Next we discuss the performance evaluations in detail.

#### 3.4.1 Gene selection for binary classes

Let us first consider the performance evaluations on binary datasets (D1-D5). In terms of Jaccard index RSGSA outperforms other algorithms in all datasets except D5. Likewise RSGSA shows better classification accuracy over other algorithms in all datasets but D4. If we take average over all datasets D1-D5, it outplays other algorithms of interest in terms of both classification accuracy and stability. Please see Table 2, Figure 3[a] and Figure 3[b] for more details. As noticeable from the performance evaluations, SVM-RFE is the closest competitor of RSGSA. Here we define % improvement as the average gain over all the datasets with respect to a performance metric. RSGSA’s % improvement over SVM-RFE ranges from 4%-7% with respect to classification accuracy. In stability measure the average % improvement over SVM-RFE ranges from 26%-48%. Please, see Figure 1[a] and Figure 1[b] for visual comparisons. To demonstrate the stability of accuracy across a set of classifiers (e.g., linear SVM (LSVM), random forest (RF), and k-nearest neighborhood (KNN)) here we took D1 dataset and performed classification accuracy over top *X* genes using 10-*fold* cross validation. It is evident from Figure 2 that all the classifiers perform equally well with respect to RSGSA’s selected top *X* genes. It further demonstrates the efficacy of our proposed method.

**Table 2.** Performance of two-class feature selection.

| Dataset | Top genes | Average Jaccard Indices (JI<sub>avg</sub>) |  |  |  |  |  | Average Informedness (I<sub>avg</sub>) |  |  |  |  |  |
| --- | --- | --- | --- | --- | --- | --- | --- | --- | --- | --- | --- | --- | --- |
|  |  | SU | GR | KLD | RELIEF | SVM-RFE | RSGSA | SU | GR | KLD | RELIEF | SVM-RFE | RSGSA |
| D1 | 50 | 0.18 | 0.21 | 0.05 | 0.29 | 0.22 | **0.38** | 0.63 | 0.62 | 0.52 | 0.67 | 0.75 | **0.86** |
|  | 100 | 0.21 | 0.21 | 0.11 | 0.31 | 0.28 | **0.41** | 0.67 | 0.68 | 0.67 | 0.64 | 0.75 | **0.85** |
|  | 150 | 0.23 | 0.23 | 0.14 | 0.32 | 0.31 | **0.44** | 0.66 | 0.67 | 0.62 | 0.62 | 0.74 | **0.84** |
|  | 200 | 0.25 | 0.25 | 0.18 | 0.33 | 0.34 | **0.47** | 0.66 | 0.68 | 0.61 | 0.62 | 0.74 | **0.86** |
| D2 | 50 | 0.04 | 0.06 | 0.02 | 0.05 | 0.14 | **0.23** | 0.14 | 0.06 | 0.05 | 0.33 | 0.56 | **0.74** |
|  | 100 | 0.05 | 0.06 | 0.03 | 0.06 | 0.19 | **0.25** | 0.21 | 0.17 | 0.01 | 0.34 | 0.59 | **0.79** |
|  | 150 | 0.07 | 0.08 | 0.05 | 0.07 | 0.21 | **0.27** | 0.19 | 0.27 | 0.11 | 0.38 | 0.63 | **0.79** |
|  | 200 | 0.08 | 0.08 | 0.07 | 0.08 | 0.24 | **0.29** | 0.18 | 0.33 | 0.16 | 0.36 | 0.68 | **0.84** |
| D3 | 50 | 0.39 | 0.44 | 0.05 | 0.24 | 0.32 | **0.56** | 0.92 | 0.90 | 0.81 | 0.93 | 0.96 | **0.99** |
|  | 100 | 0.35 | 0.43 | 0.05 | 0.22 | 0.34 | **0.52** | 0.95 | 0.92 | 0.83 | 0.92 | 0.98 | **0.99** |
|  | 150 | 0.33 | 0.43 | 0.07 | 0.22 | 0.35 | **0.53** | 0.92 | 0.92 | 0.88 | 0.91 | 0.98 | **0.99** |
|  | 200 | 0.31 | 0.42 | 0.08 | 0.22 | 0.37 | **0.53** | 0.93 | 0.93 | 0.88 | 0.94 | 0.97 | **0.99** |
| D4 | 50 | 0.04 | 0.09 | 0.01 | 0.03 | 0.12 | **0.17** | 0.38 | 0.23 | 0.26 | 0.45 | **0.55** | 0.47 |
|  | 100 | 0.05 | 0.13 | 0.02 | 0.04 | 0.17 | **0.19** | 0.39 | 0.22 | 0.31 | 0.40 | **0.57** | 0.45 |
|  | 150 | 0.06 | 0.11 | 0.03 | 0.05 | 0.18 | **0.21** | 0.38 | 0.20 | 0.25 | 0.44 | **0.61** | 0.50 |
|  | 200 | 0.07 | 0.11 | 0.04 | 0.06 | 0.20 | **0.22** | 0.40 | 0.25 | 0.29 | 0.43 | **0.59** | 0.52 |
| D5 | 50 | **0.84** | 0.78 | 0.66 | 0.61 | 0.56 | 0.65 | 0.99 | 0.99 | 0.99 | 0.99 | **1.00** | **1.00** |
|  | 100 | **0.73** | 0.65 | 0.54 | **0.73** | 0.56 | 0.63 | **1.00** | **1.00** | **1.00** | 0.99 | **1.00** | **1.00** |
|  | 150 | 0.73 | 0.61 | 0.48 | **0.81** | 0.55 | 0.64 | **1.00** | **1.00** | **1.00** | **1.00** | **1.00** | **1.00** |
|  | 200 | 0.74 | 0.60 | 0.44 | **0.81** | 0.56 | 0.65 | **1.00** | **1.00** | 0.99 | **1.00** | **1.00** | **1.00** |
| Average | 50 | 0.30 | 0.32 | 0.16 | 0.24 | 0.27 | **0.40** | 0.61 | 0.56 | 0.53 | 0.67 | 0.76 | **0.81** |
|  | 100 | 0.28 | 0.30 | 0.15 | 0.27 | 0.31 | **0.40** | 0.64 | 0.60 | 0.56 | 0.66 | 0.78 | **0.82** |
|  | 150 | 0.28 | 0.29 | 0.15 | 0.29 | 0.32 | **0.42** | 0.63 | 0.61 | 0.57 | 0.67 | 0.79 | **0.82** |
|  | 200 | 0.29 | 0.29 | 0.16 | 0.30 | 0.34 | **0.43** | 0.63 | 0.64 | 0.59 | 0.67 | 0.80 | **0.84** |
| Av. RSGSA<br>Gain over other<br>algorithms | 50 | 0.33 | 0.25 | **1.50** | 0.67 | 0.48 |  | 0.33 | 0.45 | **0.53** | 0.21 | 0.07 |  |
|  | 100 | 0.43 | 0.33 | **1.67** | 0.48 | 0.29 |  | 0.28 | 0.37 | **0.46** | 0.24 | 0.05 |  |
|  | 150 | 0.50 | 0.45 | **1.80** | 0.45 | 0.31 |  | 0.30 | 0.34 | **0.44** | 0.22 | 0.04 |  |
|  | 200 | 0.48 | 0.48 | **1.69** | 0.43 | 0.26 |  | 0.33 | 0.31 | **0.42** | 0.25 | 0.05 |  |

**Figure 1.**
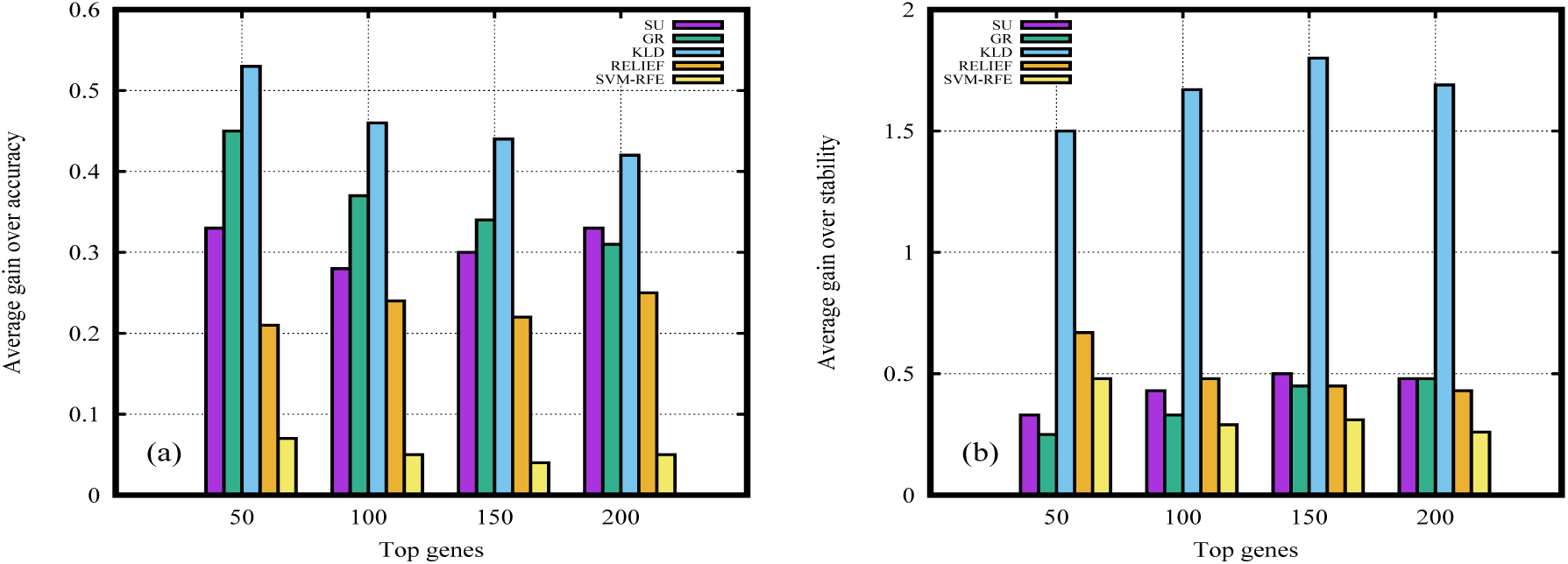
Performance evaluations of RSGSA over other notable algorithms. (a) Average gain over *informedness* (b) Average gain over *stability*

**Figure 2.**
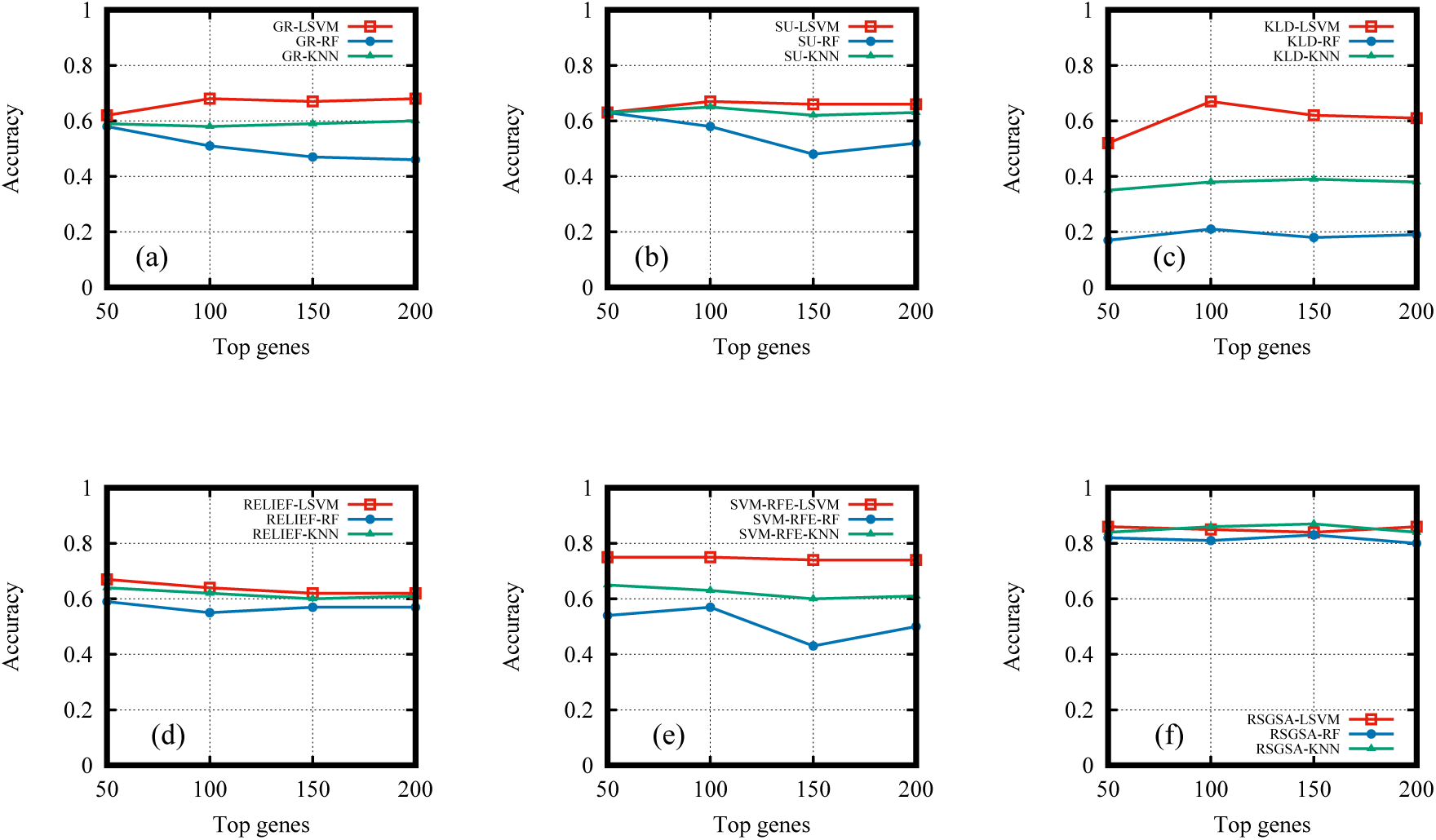
Classification accuracy of selected genes by employing LSVM, RF, and KNN classifiers for (a) GR algorithm (b) SU algorithm (c) KLD algorithm (d) RELIEF algorithm (e) SVM-RFE algorithm (f) RSGSA algorithm

**Figure 3.**
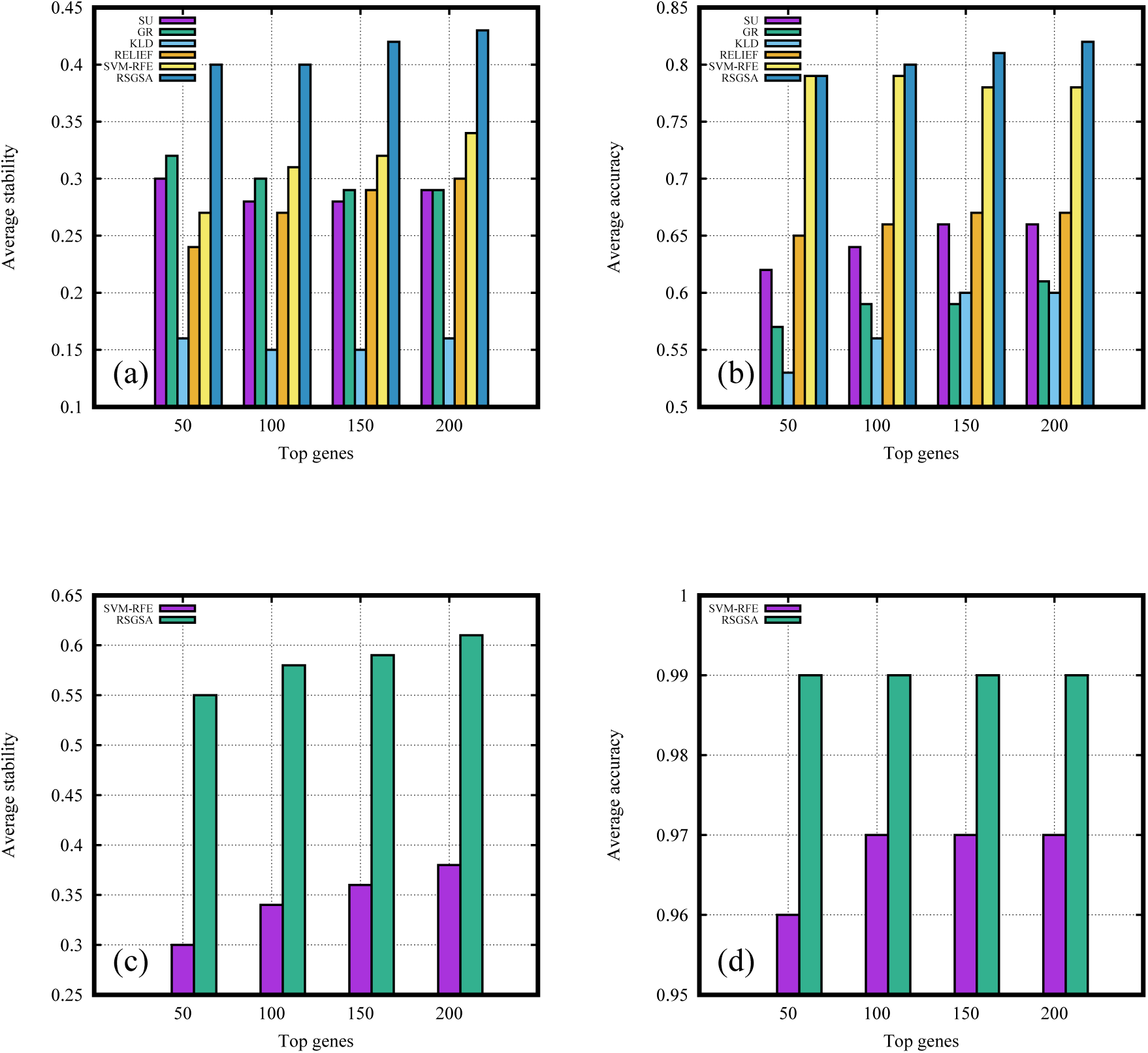
Performance evaluations (a) Average *stability* for binary class (b) Average *accuracy* for binary class (c) Average *stability* for multi class (d) Average *accuracy* for multi class

#### 3.4.2 Gene selection for multi classes

Let us now consider the performance evaluations on multi-class datasets (D6-D10). We only ran experiments by taking RSGSA and its closest competitor SVM-RFE. RSGSA outplays SVM-RFE in terms of both classification accuracy and stability for every dataset. Please see Table 3, Figure 3[c], and Figure 3[d] for more details. RSGSA’s % improvement over SVM-RFE ranges from 2%-3% with respect to classification accuracy. In stability measure the % improvement over SVM-RFE ranges from 61%-83%. Note that classification accuracy is measured by LSVM using 10-*fold* cross validation as stated above. We also performed classification accuracy based on 3 classification algorithms.

**Table 3.** Performance of multi-class feature selection.

| Dataset | Top genes | J<sub>I</sub>avg |  | I<sub>avg</sub> |  |
| --- | --- | --- | --- | --- | --- |
|  |  | SVM-RFE | RSGSA | SVM-RFE | RSGSA |
| D6 | 50 | 0.30 | **0.52** | 0.96 | **0.98** |
|  | 100 | 0.36 | **0.58** | 0.96 | **0.99** |
|  | 150 | 0.36 | **0.61** | 0.96 | **0.99** |
|  | 200 | 0.38 | **0.62** | 0.96 | **0.99** |
| D7 | 50 | 0.25 | **0.65** | 0.92 | **0.99** |
|  | 100 | 0.28 | **0.62** | 0.94 | **0.99** |
|  | 150 | 0.31 | **0.64** | 0.95 | **0.99** |
|  | 200 | 0.32 | **0.68** | 0.94 | **0.99** |
| D8 | 50 | 0.22 | **0.50** | 0.95 | **1.00** |
|  | 100 | 0.28 | **0.54** | 0.96 | **1.00** |
|  | 150 | 0.31 | **0.57** | 0.96 | **1.00** |
|  | 200 | 0.34 | **0.61** | 0.96 | **1.00** |
| D9 | 50 | 0.29 | **0.48** | **0.98** | 0.97 |
|  | 100 | 0.33 | **0.50** | 0.98 | 0.98 |
|  | 150 | 0.35 | **0.48** | 0.98 | 0.98 |
|  | 200 | 0.36 | **0.47** | 0.98 | **0.99** |
| D10 | 50 | 0.42 | **0.61** | 0.99 | **1.00** |
|  | 100 | 0.47 | **0.64** | 0.99 | **1.00** |
|  | 150 | 0.48 | **0.66** | 1.00 | 1.00 |
|  | 200 | 0.51 | **0.68** | 1.00 | 1.00 |
| Average | 50 | 0.30 | **0.55** | 0.96 | **0.99** |
|  | 100 | 0.34 | **0.58** | 0.97 | **0.99** |
|  | 150 | 0.36 | **0.59** | 0.97 | **0.99** |
|  | 200 | 0.38 | **0.61** | 0.97 | **0.99** |
| Av. Gain of<br>RSGSA over<br>SVM-RFE | 50 | 0.83 |  | 0.03 |  |
|  | 100 | 0.71 |  | 0.02 |  |
|  | 150 | 0.64 |  | 0.02 |  |
|  | 200 | 0.61 |  | 0.02 |  |

## 4 Discussion

### 4.1 Correlations

As noted in Methods section, highly correlated features normally do not improve the classification model. Next we discuss the benefits of discarding highly correlated features in detail.

#### Making the learning algorithm faster

If we have numerous correlated features it may cause the classification algorithms to consider unnecessary features while building the learning model. It eventually drives to curse of high dimensionality problem. The more the number of features the more will be execution time and memory consumption. Lower number of features generally indicates high improvement in terms of execution time and memory consumption.

#### Decreasing harmful bias

For linear models (e.g., linear regression or logistic regression), multi-collinearity can yield solutions that are wildly varying and possibly numerically unstable. Some notable models such as random forests or support vector machines can be good at detecting interactions among different features but highly correlated features can mask these interactions. This assumption can be viewed as a special case of Occam’s razor e.g., the more assumptions we have to make, the more unlikely will be an explanation.

#### Making the model simpler and interpretable

If a model of choice needs to be interpretable, the model must be simple to explain. According to Occam’s razor a simple model is desirable and a model with fewer features is simple to interpret. The concept of minimum description length makes this assumption more precise.

### 4.2 Stability and robustness

Ensuring stability and robustness mitigate 3 key issues dominating in supervised feature selection domain: (1) In a very underdetermined system where we have few hundreds to thousands of samples with thousands to million of features (e.g., DNA microarray, RNA-seq data, GWAS data) it is often found that contrasting feature subsets of same size may yield equally identical results [31]. Aggregating a set of feature selection algorithms can aid to reduce the risk of erroneously favoring an unstable subset of features; (2) Different feature selection methods may fall in local optima in the space of feature subsets. Therefore, individually each of the algorithms can produce unstable results. Ensembling multiple feature selection techniques may better cater approximation to the optimal subset or ranking of features; (3) Finally the hypothesis space searched by an algorithm alone might not contain the true target function. By acting together in a concerted way a set of classifier can produce good approximation over the hypothesis space.

As noted in Methods section, we attempt to make our learning model stable by introducing a small amount of random “noise” in the training examples. E.g., in each recursive step of SSVM-RFE each linear SVM from the set of ensembles is trained with slightly different examples by randomly flipping class labels of few examples. Like any other learning algorithm SVM is stochastic in nature and so, it is very sensitive with the training examples. As a consequence each SVM will produce slightly different hyperplane and we will end up with different values in weight associated with each gene. The claim is if a gene is stable (or, discriminating) it will remain stable within small “noise.” At the end, SSVM-REF assigns ranks to every gene in the dataset. The lower the rank of a particular gene the higher will be its importance. To ensure robustness we bootstrap the training examples multiple times i.e., we randomly select a subset of training examples from entire set of training examples with replacement multiple times. For each bootstrapped samples we execute SSVM-REF and get ranks based on their importance for all the genes. Finally, we aggregate gene ranks produced by each SSVM-REE by using Equation 3.

To demonstrate the effectiveness of RSGSA in terms of *stability*, we show outcome of D1 dataset in Figure 4 by ensembling 5 linear SVMs in one recursive stage of SSVM-RFE. One of the notable statistical measures of *stability* is coefficient of variation (CV). It is a standardized measure of dispersion of a probability distribution or frequency distribution. CV is defined as the ratio of the standard deviation *σ* to the mean *µ* [8]: 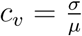 It is generally expressed as a percentage. The higher the coefficient of variation, the greater will be the level of dispersion around the mean. Therefore, the lower the value of the coefficient of variation, the more precise will be the estimate. Consider 10 most important and 10 least important genes according to their weight distribution coming from 10 linear SVMs. CVs of 10 most important genes varies from 4% to 15% (please see, Figure 4[b]). Boxplots in Figure 4[a] also support the evidence. On the contrary for 10 least important genes, it varies from 60% to 170% (please see, Figure 4[d]). Boxplots in Figure 4[c] also support the findings. Figure 4[e] shows the weights of every gene in D1 dataset produced by 5 linear SVMs in the first recursive stage of SSVM-RFE. Please note that we randomly introduced “noise” by flipping 3% class labels before executing each linear SVM.

**Figure 4.**
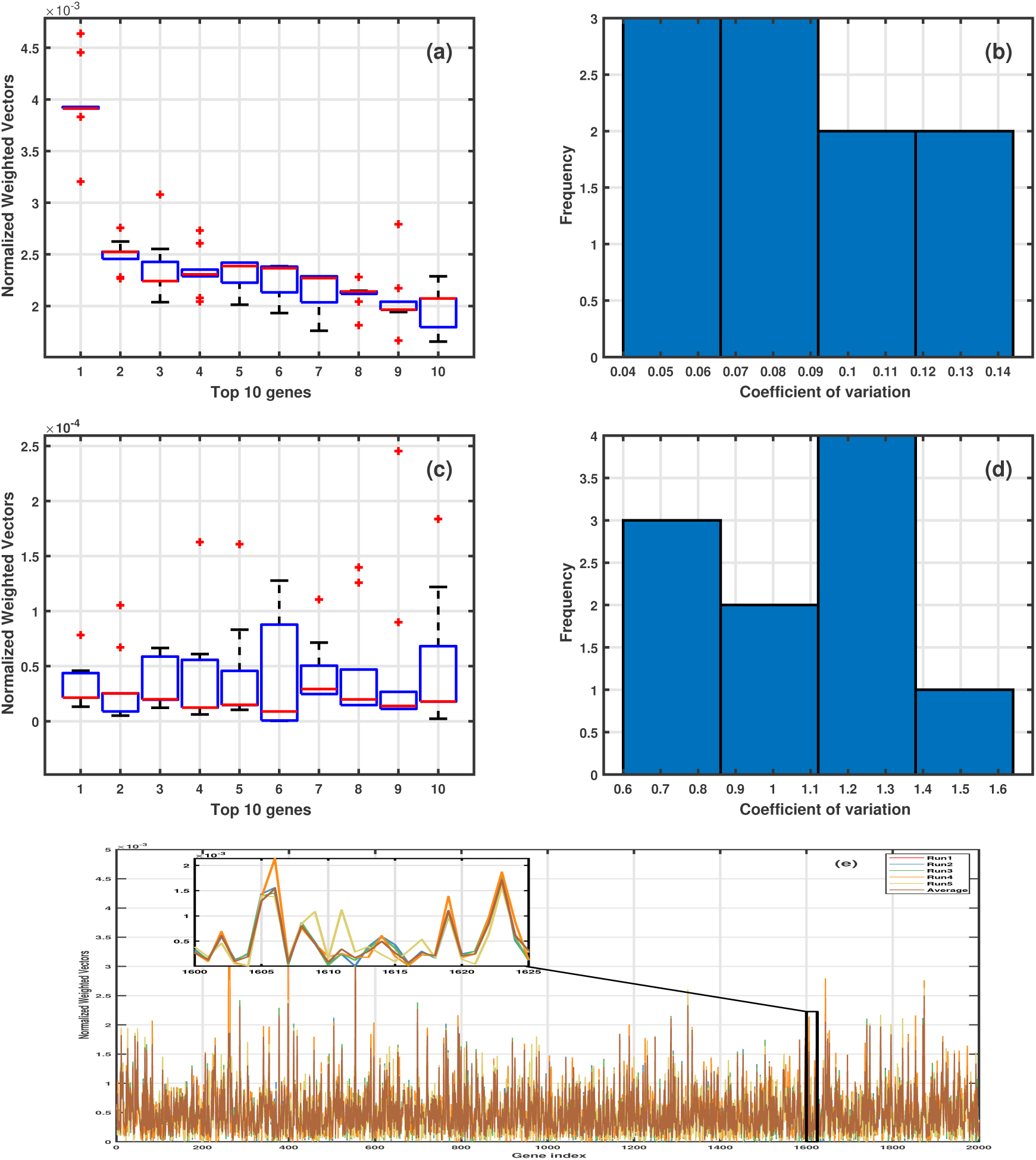
Outcome of a recursive stage of SSVM-RFE (a) Weights of top 10 genes (b) CV of top 10 genes (c) Weights of least 10 genes (d) CV of least 10 genes (e) Weights of all genes produced by 5 LSVM and their average in a recursive stage

### 4.3 Biological significance

We have enlisted top 10 most important genes in Table 4. They have been selected from D1 dataset by our proposed algorithm RSGSA. Next we briefly discuss our findings considering the top 3 genes. T-cell-specific transcription factor 7 (TCF7) is directly associated with colorectal cancer [40]. The second one, TPM4, is associated with clinical progression in colon cancer patients and acts as a tumor suppressor in colon cancer cells [https://www.ncbi.nlm.nih.gov/gene/7171]. Finally, ACTB is closely associated with a variety of cancers and accumulating evidence indicates that ACTB is de-regulated in liver, melanoma, renal, colorectal, gastric, pancreatic, esophageal, lung, breast, prostate, ovarian cancers, leukemia and lymphoma [12].

**Table 4.** Top 10 genes selected by RSGSA from D1 dataset.

| Rank | Probe ID | Gene symbol | Full name |
| --- | --- | --- | --- |
| 0 | X59871 | TCF7 | transcription factor 7 |
| 1 | X05276 | TPM4 | tropomyosin 4 |
| 2 | X63432 | ACTB | actin beta |
| 3 | J05032 | DARS1 | aspartyl-tRNA synthetase 1 |
| 4 | D26535 | DLST | dihydrolipoamide S-succinyltransferase |
| 5 | H68220 | FAU | FAU ubiquitin like and ribosomal protein S30 fusion |
| 6 | T97199 | ITGB4 | integrin subunit beta 4 |
| 7 | T56244 | PSMB2 | proteasome subunit beta 2 |
| 8 | R16255 | PPP3CB | protein phosphatase 3 catalytic subunit beta |
| 9 | T70063 | EIF4G2 | eukaryotic translation initiation factor 4 gamma 2 |

Now consider the biological significance of the top 100 genes selected from colon tumor dataset (D1) by our proposed method RSGSA. The enrichment analysis is based on three dimensions e.g., gene ontology-biological processes (GO-BP) terms, disease ontology (DO) terms, and biological pathways. Please, note that GO-BP and DO analyses were performed using “clusterProfiler” [42]. Biological pathways are extracted from “ConsensusPathDB-human” (CPDB, in short) [http://cpdb.molgen.mpg.de/]. Next we briefly illustrate each of the enrichment analysis.

#### Gene Ontology-Biological Processes (GO-BP)

One of the main uses of the GO terms is to perform enrichment analysis on a given set of genes. For instance, an enrichment analysis will find which GO terms are over-represented (or under-represented) using annotations for that set of genes. We have performed enrichment analysis on the set of 100 genes as noted above based on GO-BP terms and retained 104 GO-BP Benjamini-Hochberg corrected (adjusted *p* < 0.05) terms. Top 10 enriched GO-BP terms can be found in Table 5. Most of the terms are associated with colon cancer. For example the authors in [37] have described CRC and Renal Cell Carcinoma (RCC) as concomitant malignancies in their study of patients carrying both types of cancer. Patients with Retinoblastoma are also shown to have high risk of getting colon cancer over time [19].

**Table 5.**
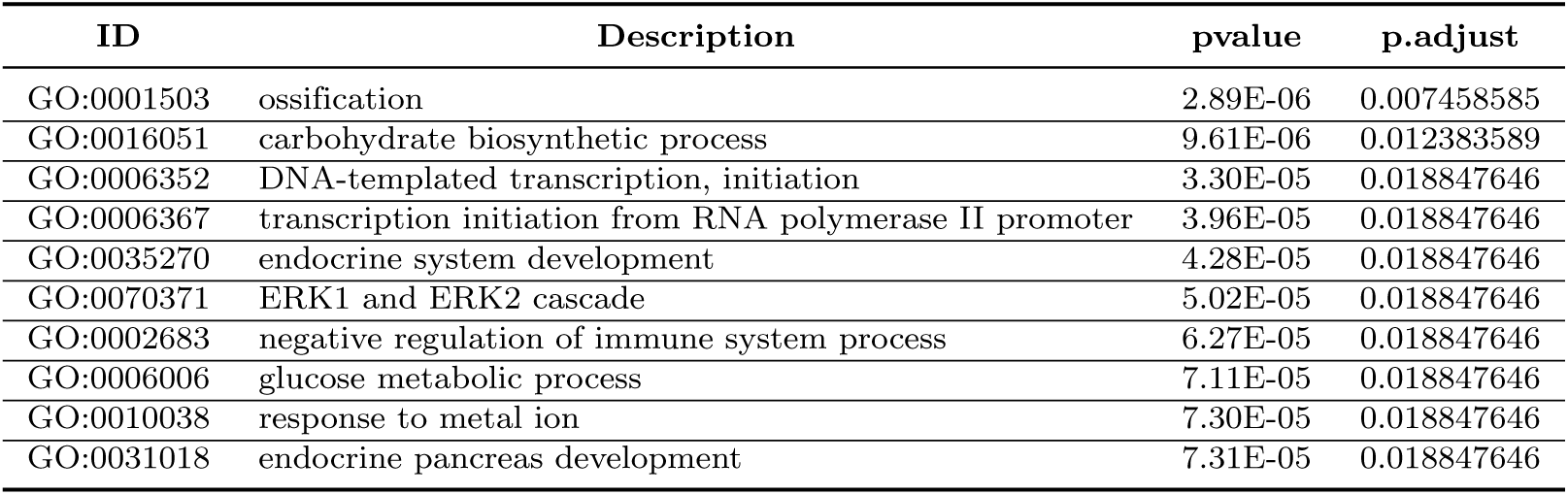
Top 10 enriched (GO-BP) terms.

#### Disease Ontology (DO)

Like GO, the disease ontology (DO) is a formal ontology of human disease. We have performed enrichment analysis on the set of top 100 genes as noted above based on DO terms and retained 17 DO Benjamini-Hochberg corrected (adjusted *p* < 0.05) terms. Top 10 enriched DO terms can be found in Table 6. Almost all of the retained enriched DO terms are associated with colon cancer. For instance although a rare case, it has been reported that a 76-year-old woman has a colon cancer with ossification [26]. For another instance it has been shown that colon cancer progression has been impaired via inactivating the Wnt pathway [41].

**Table 6.** Top 10 enriched disease ontology (DO) terms.

| ID | Description | p-value | p.adjust |
| --- | --- | --- | --- |
| DOID:3996 | urinary system cancer | 0.000161338 | 0.031918678 |
| DOID:4450 | renal cell carcinoma | 0.00027404 | 0.031918678 |
| DOID:0060116 | sensory system cancer | 0.000278766 | 0.031918678 |
| DOID:2174 | ocular cancer | 0.000278766 | 0.031918678 |
| DOID:4451 | renal carcinoma | 0.000670596 | 0.03438733 |
| DOID:768 | retinoblastoma | 0.000674808 | 0.03438733 |
| DOID:771 | retinal cell cancer | 0.000674808 | 0.03438733 |
| DOID:4645 | retinal cancer | 0.000738581 | 0.03438733 |
| DOID:14067 | plasmodium falciparum malaria | 0.000771616 | 0.03438733 |
| DOID:2377 | multiple sclerosis | 0.000903255 | 0.03438733 |

#### Biological Pathways

We have also performed biological pathway analysis and retained 23 Bonferroni adjusted (*p* < 0.05) enriched pathways. Almost all of the retained pathways are associated with colon cancer. Top 10 enriched biological pathways can be found in Table 7. Now we illustrate some of the pathways in detail. Interleukins are a group of cytokines that contribute to growth and differentiation, cell migration, and inflammatory and anti-inflammatory responses by the immune system. In a study, authors in [6] examined genetic variation in genes from various anti-inflammatory and pro-inflammatory interleukins to determine association with colon and rectal cancer risk and overall survival.

**Table 7.** Top 10 enriched biological pathways.

| Pathway | Source | ID | Hypergeometric p-value |
| --- | --- | --- | --- |
| IL3 | NetPath | Pathway_IL3 | 2.75E-06 |
| Cellular senescence | KEGG | path:hsa04218 | 4.51E-06 |
| CD4 T cell receptor signaling | INOH | None | 8.12E-06 |
| VEGF | INOH | None | 1.59E-05 |
| Alpha6Beta4Integrin | NetPath | Pathway_Alpha6Beta4Integrin | 1.93E-05 |
| TCR | NetPath | Pathway_TCR | 2.14E-05 |
| Fibroblast growth factor-1 | NetPath | Pathway_Fibroblast__growth__factor-1 | 2.44E-05 |
| a6b1 and a6b4 Integrin signaling | PID | a6b1_a6b4_integrin_pathway | 2.59E-05 |
| BCR | NetPath | Pathway_BCR | 6.87E-05 |
| EPO signaling | INOH | None | 1.02E-04 |

Data from two population-based incident studies of colon cancer (1,555 cases and 1,956 controls) and rectal cancer (754 cases and 954 controls) were utilized. After controlling for multiple comparisons, authors found that single nucleotide polymorphisms (SNPs) from four genes, IL3, IL6R, IL8, IL15, were associated with increased colon cancer risk. It has also been discovered that colorectal cancer cells express a hybrid form of *α*6*β*4 that is never seen in normal cells [4]. The expression levels of epidermal growth factor receptors (EGFRs) varies significantly on normal and malignant colon epithelial cells [33]. Furthermore, activation of the EGFR signaling pathway was proposed as a rational target for anti-tumor drugs [2]. Human T-lymphotropic virus-I (HTLV-I) is one of the retroviruses associated with human cancer [24].

## 5 Conclusion

In this article we have proposed a robust and stable supervised gene selection algorithm RSGSA based on graph theory and ensembles of linear SVMs. At the beginning, highly correlated genes are discarded by employing a novel graph theoretic algorithm. Stability of SVM-RFE is ensured by introducing small “noise” in phenotypes (i.e., class labels). Robustness is secured by instance level perturbation (i.e, bootstrapping samples multiple times). Rigorous experimental evaluations were performed on 10 real gene expression datasets. It is indisputable from the evaluations that RSGSA is indeed an effective and efficient supervised gene selection algorithm.

## Availability of data and materials

The implementations and the datasets we used are freely available for non-commercial purposes. They can be downloaded from: https://figshare.com/projects/RSGSA_Robust_and_Stable_Gene_Selection_Algorithm/67634

## References

1. J. Alles, T. Fehlmann, U. Fischer, C. Backes, V. Galata, M. Minet, M. Hart, M. Abu-Halima, F. A. Grässer, H.-P. Lenhof, A. Keller, and E. Meese. An estimate of the total number of true human miRNAs. Nucleic Acids Research, 47(7):3353–3364, Apr. 2019.

2. C. L. Arteaga. Epidermal Growth Factor Receptor Dependence in Human Tumors: More Than Just Expression? The Oncologist, 7(Supplement 4):31–39, Aug. 2002.

3. D. J. Balding. A tutorial on statistical methods for population association studies. Nature Reviews Genetics, 7(10):781–791, Oct. 2006.

4. J.-F. Beaulieu. Integrin α6β4 in colorectal cancer. World Journal of Gastrointestinal Pathophysiology, 1(1):3–11, Apr. 2010.

5. V. Bolón-Canedo and A. Alonso-Betanzos. Ensembles for feature selection: A review and future trends. Information Fusion, 52:1–12, Dec. 2019.

6. K. L. Bondurant, A. Lundgreen, J. S. Herrick, S. Kadlubar, R. K. Wolff, and M. L. Slattery. Interleukin genes and associations with colon and rectal cancer risk and overall survival. International Journal of Cancer, 132(4):905–915, Feb. 2013.

7. L. Breiman. Random Forests. Machine Learning, 45(1):5–32, Oct. 2001.

8. B.. Brian S. Everitt. Cambridge Dictionary of Statistics. Cambridge University Press, 1 edition, Oct. 1998.

9. K. Bryan and T. Leise. The $25,000,000,000 Eigenvector: The Linear Algebra behind Google. SIAM Review, 48(3):569–581, Jan. 2006.

10. N. V. Chawla, K. W. Bowyer, L. O. Hall, and W. P. Kegelmeyer. SMOTE: Synthetic Minority Over-sampling Technique. Journal of Artificial Intelligence Research, 16(1):321–357, June 2002. 1106.1813.

11. I. Ezkurdia, D. Juan, J. M. Rodriguez, A. Frankish, M. Diekhans, J. Harrow, J. Vazquez, A. Valencia, and M. L. Tress. Multiple evidence strands suggest that there may be as few as 19,000 human protein-coding genes. Human Molecular Genetics, 23(22):5866–5878, Nov. 2014.

12. C. Guo, S. Liu, J. Wang, M.-Z. Sun, and F. T. Greenaway. ACTB in cancer. Clinica Chimica Acta; International Journal of Clinical Chemistry, 417:39–44, Feb. 2013.

13. I. Guyon, J. Weston, S. Barnhill, and V. Vapnik. Gene Selection for Cancer Classification using Support Vector Machines. Machine Learning, 46(1-3):389–422, Jan. 2002.

14. L. K. Hansen and P. Salamon. Neural network ensembles. IEEE Transactions on Pattern Analysis and Machine Intelligence, 12(10):993–1001, Oct. 1990.

15. Z. Hu, H. S. Scott, G. Qin, G. Zheng, X. Chu, L. Xie, D. L. Adelson, B. E. Oftedal, P. Venugopal, M. Babic, C. N. Hahn, B. Zhang, X. Wang, N. Li, and C. Wei. Revealing Missing Human Protein Isoforms Based on Ab Initio Prediction, RNA-seq and Proteomics. Scientific Reports, 5:15, July 2015.

16. International Human Genome Sequencing Consortium. Finishing the euchromatic sequence of the human genome. Nature, 431(7011):931–945, Oct. 2004.

17. A. G. Karegowda, A. S. Manjunath, and M. A. Jayaram. COMPARATIVE STUDY OF ATTRIBUTE SELECTION USING GAIN RATIO AND CORRELATION BASED FEATURE SELECTION. International Journal of Information Technology and Knowledge Management, 2(2):271–277, Dec. 2010.

18. S. S. Keerthi and D. DeCoste. A Modified Finite Newton Method for Fast Solution of Large Scale Linear SVMs. J. Mach. Learn. Res., 6:341–361, 2005.

19. R. A. Kleinerman, M. A. Tucker, R. E. Tarone, D. H. Abramson, J. M. Seddon, M. Stovall, F. P. Li, and J. F. Fraumeni. Risk of new cancers after radiotherapy in long-term survivors of retinoblastoma: an extended follow-up. J Clin Oncol, 2005.

20. R. Kohavi and G. H. John. Wrappers for feature subset selection. Artificial Intelligence, 97(1):273–324, Dec. 1997.

21. R. Kohavi and G. H. John. Wrappers for feature subset selection. Artificial Intelligence, 97(1):273–324, Dec. 1997.

22. I. Kononenko. Estimating attributes: Analysis and extensions of RELIEF. In F. Bergadano and L. De Raedt, editors, Machine Learning: ECML-94, Lecture Notes in Computer Science, pages 171–182. Springer Berlin Heidelberg, 1994.

23. R. Meiri and J. Zahavi. Using simulated annealing to optimize the feature selection problem in marketing applications. European Journal of Operational Research, 171(3):842–858, June 2006.

24. P. S. Moore and Y. Chang. Why do viruses cause cancer? Highlights of the first century of human tumour virology. Nature Reviews Cancer, 10(12):878–889, Dec. 2010.

25. C. F. A. Negre, U. N. Morzan, H. P. Hendrickson, R. Pal, G. P. Lisi, J. P. Loria, Rivalta, J. Ho, and V. S. Batista. Eigenvector centrality for characterization of protein allosteric pathways. Proceedings of the National Academy of Sciences, 115(52):E12201–E12208, Dec. 2018.

26. B.-J. Noh, Y. W. Kim, and Y.-K. Park. A Rare Colon Cancer with Ossification: Pathogenetic Analysis of Bone Formation. Annals of Clinical and Laboratory Science, 46(4):428–432, July 2016.

27. V. Y. Pan and Z. Q. Chen. The Complexity of the Matrix Eigenproblem. In Proceedings of the Thirty-first Annual ACM Symposium on Theory of Computing, STOC ‘99, pages 507–516, New York, NY, USA, 1999. ACM. event-place: Atlanta, Georgia, USA.

28. V. Y. Pan, Z. Q. Chen, and A. Zheng. The Complexity of the Algebraic Eigenproblem. Mathematical Sciences Research Institute, 1998.

29. W. H. Press, S. A. Teukolsky, W. T. Vetterling, and B. P. Flannery. Numerical Recipes in C: The Art of Scientific Computing. Second Edition. Cambridge University Press, 1992.

30. T. H. Saey. A recount of human genes ups the number to at least 46,831, Sept. 2018.

31. Y. Saeys, I. Inza, and P. Larrañaga. A review of feature selection techniques in bioinformatics. Bioinformatics, 23(19):2507–2517, Oct. 2007.

32. S. Saha, S. Rajasekaran, and R. Ramprasad. Novel Randomized Feature Selection Algorithms. International Journal of Foundations of Computer Science, 26(03):321–341, Apr. 2015.

33. D. S. Salomon, R. Brandt, F. Ciardiello, and N. Normanno. Epidermal growth factor-related peptides and their receptors in human malignancies. Critical Reviews in Oncology/Hematology, 19(3):183–232, July 1995.

34. F. Santosa and W. Symes. Linear Inversion of Band-Limited Reflection Seismograms. SIAM Journal on Scientific and Statistical Computing, 7(4):1307–1330, Oct. 1986.

35. R. E. Schapire. The strength of weak learnability. Machine Learning, 5(2):197–227, June 1990.

36. S. Shedbale and D. K. Shaw. N-Gram and KLD Based Efficient Feature Selection Approach for Text Categorization. International Journal of Advance Engineering and Research Development, 4(6):9, June 2017.

37. E. Steinhagen, H. G. Moore, S. A. Lee-Kong, J. Shia, A. Eaton, A. J. Markowitz, P. Russo, and J. G. Guillem. Patients With Colorectal and Renal Cell Carcinoma Diagnoses Appear to Be at Risk for Additional Malignancies. Clinical Colorectal Cancer, 12(1):23–27, Mar. 2013.

38. A. N. Tikhonov and V. I. Arsenin. Solutions of ill-posed problems. Winston, 1977. Google-Books-ID: ECrvAAAAMAAJ.

39. L. Vermeulen-Jourdan, C. Dhaenens, and E.-G. Talbi. Linkage disequilibrium study with a parallel adaptive ga. International Journal of Foundations of Computer Science, 16(02):241–260, Apr. 2005.

40. M. L. Waterman. Lymphoid enhancer factor/T cell factor expression in colorectal cancer. Cancer Metastasis Reviews, 23(1-2):41–52, June 2004.

41. D. Yan, W. Liu, Y. Liu, and M. Luo. LINC00261 suppresses human colon cancer progression via sponging miR-324-3p and inactivating the Wnt/β-catenin pathway. Journal of cellular physiology, June 2019.

42. G. Yu, L.-G. Wang, Y. Han, and Q.-Y. He. clusterProfiler: an R package for comparing biological themes among gene clusters. Omics: A Journal of Integrative Biology, 16(5):284–287, May 2012.

43. Z. Zhu, Y.-S. Ong, and M. Dash. Markov blanket-embedded genetic algorithm for gene selection. Pattern Recognition, 40(11):3236–3248, Nov. 2007.

